# Pre-clinical in vitro and in vivo characterization of a maternal vaccination before conception to protect against severe neonatal infections caused by *Escherichia coli* K1

**DOI:** 10.1101/2022.12.29.522168

**Authors:** Youssouf Sereme, Cécile Schrimp, Esther Lefebvre-Wloszczowski, Maeva Agapoff, Helène Faury, Yunhua Chang Marchand, Elisabeth Agiron-Ardila, Emilie Panafieu, Frank Blec, Mathieu Coureuil, Eric Frappy, Stephane Bonacorsi, David Skurnik

## Abstract

Preterm birth remains the leading cause of neonatal morbidity and mortality today. Genetic, immunological, and infectious substrates are suspected. Preterm infants are at higher risk of severe neonatal infections and the main cause of bacterial infection in this population is *Escherichia coli* K1. Unfortunately, women with history of preterm birth have a high risk of recurrence. Therefore, these women constitute a target population for a vaccine, to date non-existent, against *E. coli* K1 to prevent these infections.

In this study, we characterize the immunological and microbiological properties in adult female mice of a live attenuated vaccine candidate and the protection it conferred to newborn mice against severe infection caused by *E. coli* K1. We show that our *E. coli* K1 Δ*aro*A vaccine induces a strong immunity driven by polyclonal bactericidal antibodies. In our model of meningitis, pups born from mothers immunized before conception were strongly protected against different strains of *E. coli* K1 both in early-onset and late-onset diseases.

Given the very high rate of mortality and neurological sequalae in neonatal meningitis caused by *E. coli* K1, this pre-clinical study provides a proof-of-concept for the development of a vaccine strategy against *E. coli* K1 severe infection in women at risk of preterm birth.

## Introduction

Defined by the World Health Organization (WHO) as a birth occurring before 37 completed weeks of gestation, preterm birth remains the leading cause of neonatal morbidity and mortality today^1^. Each year, approximately 15 million babies are born prematurely worldwide, representing a rate of 11% of births, with 90% of these preterm births occurring in low- and resource-poor countries^2^. With a death rate of 35% for newborns and 18% for children under 5, prematurity remains a real global public health problem^2,3^. The etiology of preterm birth is multifactorial, among which a genetic, infectious and immunological substrates are suspected^4,5^. Studies have described that women with an history of a first preterm delivery were significantly more likely to have a second preterm birth^5^.

Preterm infants have higher predisposing factors for neonatal infections compared to full-term infants due to their impaired immune system^6^. Indeed, preterm infants have functional impairment of innate immunity, decreased cytokines, chemokines, antimicrobial proteins and peptides and the complement system and immature cellular immunity^6–9^. *E. coli* is one of the most common pathogens cause of neonatal infections in preterm and term infants and the leading cause of neonatal bacterial meningitis in preterm infants^10–12^. Studies have reported significantly higher rates of vaginal *E. coli* carriage in women with preterm births compared to term controls^7^ and 90% of Gram-negative isolates identified in women with imminent preterm birth were E. coli^13,14^. Women who have had a preterm birth are a population whose future newborns will be at high risk for neonatal *E. coli K1* meningitis and therefore, vaccine prevention in women with a premature first birth could significantly reduce neonatal meningitis.

Scientific and technological progress have been made towards the development of a vaccine against *E. coli* K1 bacterial meningitis and while for example a vaccine candidate against *E. coli* K1 protein based on the OmpA structure has shown efficacy in mice^8,15,16^, to date there is no vaccine clinically available against *E. coli* K1.

Live attenuated vaccines have been reported to confer the high level of rapid and sustained full immunity necessary for maternal and offspring protection^17,18^. Although they are highly effective, the risk of reversion to a more virulent strain of the pathogen against which vaccination is given cannot be ruled out, although it is minor. Therefore, the use of these vaccines is not recommended in certain venerable populations such as pregnant women and immunocompromised individuals^19,20^. However, no contraindication to live vaccines has been reported for young women of childbearing age considering pregnancy or for women at risk of preterm birth^20–22^. To date, no live attenuated vaccine for neonatal protection against *E. coli K*1 meningitis has been reported.

Using saturated transposon (Tn) mutagenesis and high-throughput sequencing (TnSeq), a powerful tool to study host-pathogen interactions, we recently identified the *aro*A gene as a virulence factor for *E. coli* K1^23^. It is a gene that encodes 5-enopyruvylshikimate-3-phosphate synthase (EPSPS) and is involved in the biosynthesis of aromatic amino acids^24^. Live vaccines attenuated by deletion of the *aro*A gene have shown strong protection against *Pseudomonas aeruginosa* and *Salmonella*^25,26^. The efficacy of an attenuated *E. coli* O78 vaccine by deletion of the *aro*A gene against avian diseases has also been reported^27^.

In this study, after confirming that *aro*A was a virulence factor for *E. coli* K1, we characterize the immunological and microbiological properties of a live attenuated *E. coli K1* Δ*aro*A vaccine candidate injected in adult female mice. We then demonstrate the very strong protection of infant mice born from mothers immunized with this attenuated vaccine before gestation. Our results present a preclinical proof-of-concept new strategy for vaccine development against *E. coli* K1 for women at risk of premature birth.

## Results

### Characterisation and evaluation of the virulence of the aroA gene deletion E. coli K1 strain

#### *Construction of* E. coli *K1 ΔaroA*

Using the methodology described by Datsenko^28^, we constructed a *∆aroA* mutant in *E. coli* E11^23^. The deletion of the *aroA* gene in the mutant strain was confirmed by PCR analysis in the presence of controls. Clones resistant to 100 ug/mL kanamycin were obtained and were tested by PCR to verify the insertion of the kanamycin cassette and the deletion of the *aroA* gene (**Figure 1A**).

**Figure 1:**
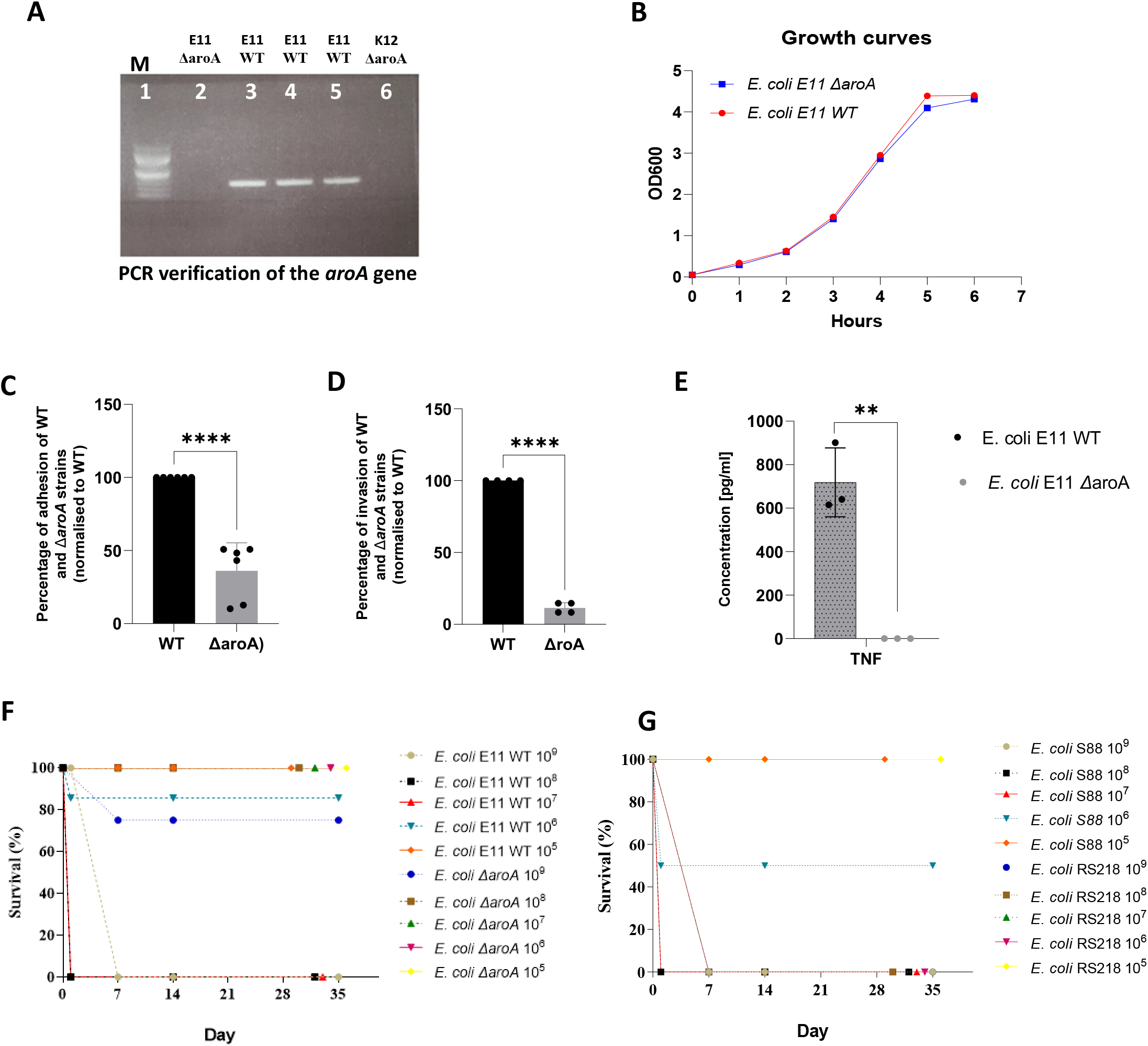
Characterization and evaluation of the virulence of the *E. coli K1* ΔaroA strain Verification PCR image of the production of the *E. coli* E11 aroA deletion mutant. A partial genetic sequence of aroA was replaced by the kanamycin resistant cassette at the cleavage sites. Identification of the aroA *E. coli* E11 mutant (**A**). Pathway 1: DNA marker. Lane 2: No PCR product for the mutant strain using the aroA-Fwd/aroA-Rev primer pair. Lane 3, 4 and 5 positive control *E. coli* E11 WT. Lane 5: Negative control *E. coli* K12 aroA.Growth curves of *E. coli* E11 WT and ΔaroA strains in LB (**B**). After 7 h of culture, there was no significant difference in the growth rate of the two strains grown in LB broth (p < 0.05). Adhesion to (**C**), invasion (**D**) of *Hela* cells by *E. coli E11 Δ*aroA were significantly reduced, compared to the wild-type strain *E. coli E11*. E11 ΔaroA significantly reduced adhesion and invasion by 25.38% and 11.49% (p<0.05). **(E)** TNF secretion by Hela cells during invasion by *E. coli E11 Δ*aroA was significantly reduced, compared to the wild-type E11 strain (p<0.05). Evaluation of in vivo mortality of the mutant strain compared to the virulent wild-type *E. coli* E11, S88 and RS218 strains. The wild-type strains were 1000 times more virulent than the mutant strain (**Figure 1F and 1G)**. The lethal dose of the virulent strains was 107 for *E. coli* E11 WT and S88, 106 for RS218 and over 109 for the mutant strain.

#### *In vitro and in vivo evaluation of aroA as a virulence factor in* E. coli *K1*

The growth rate of the *aro*A-deleted strain was similar to that of the wild-type *E. coli* E11 WT strain after seven hours of culture in LB medium (**Figure 1B**). However, comparing the adhesion and invasion capacity of the mutant strain to the wild-type strain, we found that the adhesion and invasion of the *aroA* gene-deleted strain on Hela epithelial cells was significantly reduced compared to the wild-type strain. The p-values were less than 0.0001 (**Figure 1C and D)**. In addition, epithelial cells stimulated for four days with the deletion strain did not secrete TNFα in contrast to the WT strain (**Figure 1E**).

When evaluating the in vivo mortality by the mutant strain E11 Δ*aro*A compared to the wild-type strain E11 and two other virulent strains (*E. coli* S88^29^ and RS218^30^), we found that the wild-type E11 strain and the other two virulent strains were 1000-fold more virulent than the mutant strain. The lethal doses were 10^7^ for *E. coli* E11 WT and *E. coli* S88, 10^6^ for *E .coli* RS218 and over 10^9^ for the mutant strain E11 Δ*aro*A (**Figure 1F** and **1G)**

### Evaluation of the humoral response after immunization

Having confirmed that *aro*A was a virulence factor in *E. coli* K1 and that the strain E11 Δ*aro*A could be used as a live attenuated vaccine, we next evaluated the immunogenicity of this vaccine candidate. Following the experimental immunization design presented **Figure. 2A**, after intraperitoneal inoculation with 10^5^ CFU of *E. coli* E11 Δ*aro*A bacteria, vaccinated female mice were sampled for humoral response testing. Antibody levels specific to three *E. coli* strains responsible for neonatal meningitis (E11, S88 and RS218)^29,30^ were detected in the sera of immunized mice (**Figure 2B**) and a sharp increase in levels was observed after the third dose (**Figure 2C**). When testing specific antibodies to other K1 and non-K1 *E. coli* strains responsible of clinical neonatal meningitis an increased level of antibodies was found in the sera of immunized mice compared to sera from the pre-immune mice (**Supplementary figure 1A and 1B**). To confirm that the binding to several strains of *E. coli* K1 was caused by a polyclonal response, we next performed an SDS PAGE gel and Western Blots with three *E. coli* K1 strains (**Figure 2F-H**). We were able to visualize the proteins of these bacteria (**Figure 2F**) and to detect polyclonal antibodies against several antigens of these bacteria by western blot (**Figure 2G-H**). Together, these results show that immunization with the *aro*A mutant *E. coli* K1 strain induced an humoral response with polyclonal antibodies able to bind numerous different strains of *E. coli* K1.

**Figure 2:**
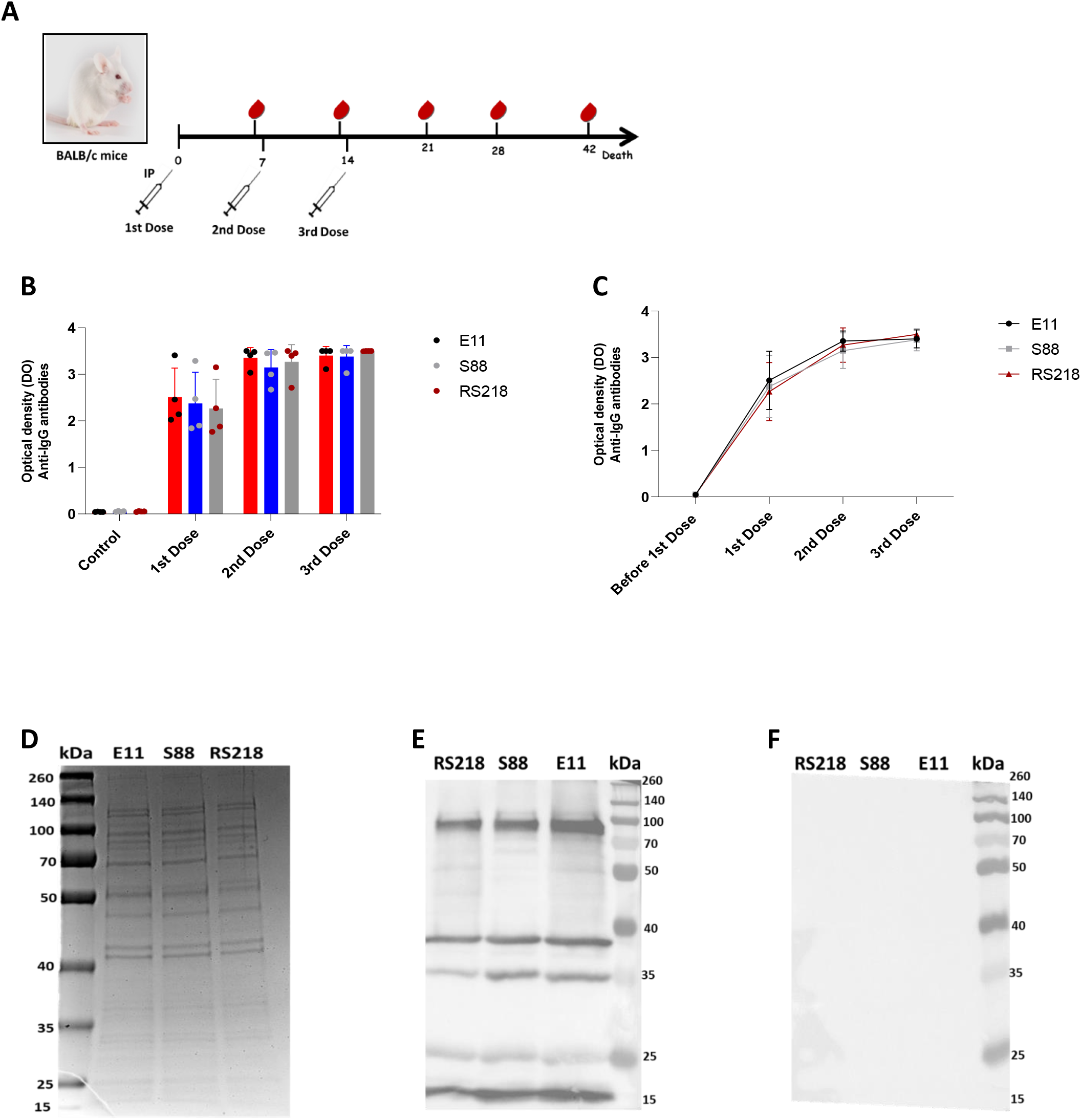
Evaluation of the humoral response after immunization by *E. coli* E11 ΔaroA **(A)** Schema of immunization of Balb/C female mice with 105 CFU of *E. coli K1* E11 ΔaroA (n=4) or NaCl sham (n=4=). (**B**) IgG response to the three major *E*.*coli K1* strains (E11, S88 and RS218) responsible for neonatal meningitis after each dose. (**C**) IgG response kinetics to the *three E. coli* strains E11, S88 and RS218 before immunisation and after each dose. **(D)** SDS-PAGE analyses with Coomassie Blue staining for *E. coli* E11, S88 and RS218 antigens. In the profile, the left lane corresponds to the antigen lane and the left lane corresponds to the molecular weight standard and; 6 µg of the protein extracts of the three strains were run in a 12% polyacrylamide gel *E. coli E11*, S88 and RS218. **(E** and **F**) Western blot of *E. coli* antigens, detection of anti-mouse IgG antibodies was performed in the serum of one immunized mouse (**E**) and control (non-immunised) serum (**F**).

### Evaluation of the cellular response after immunization

We next evaluated the cellular response in female Balb/C mice immunized with three doses of E11 Δ*aro*A. Seven days after the third dose of immunization, mice were sacrificed to collect the spleens, and splenocytes were isolated and analyzed. Sera were analyzed to quantify cytokines of the Th1, Th2 and Th17 response. The results showed that there was no significant difference (Student test, p>0.05) in the expression of T cell (**Figure 3A**), B cell (**Figure 3B**), CD4 and CD8 T cell populations (**Figure 3C and 3D**) and regulatory T cells (**Figure 3 E and F**) in the groups of vaccinated mice compared to the unvaccinated animals. However, a trend of increasing the regulatory T population was observed in splenocytes of the vaccinated mice (**Figure 3E**). At the systemic level, the immunized mice induced more TNFα (**Figure 3I**) and IL-10 (**Figure 3G**) compared to the non-immune groups even though the level of these detected cytokines was very low. These results indicate that vaccination with the E11 Δ*aro*A strain does not induce a robust cellular response.

**Figure 3:**
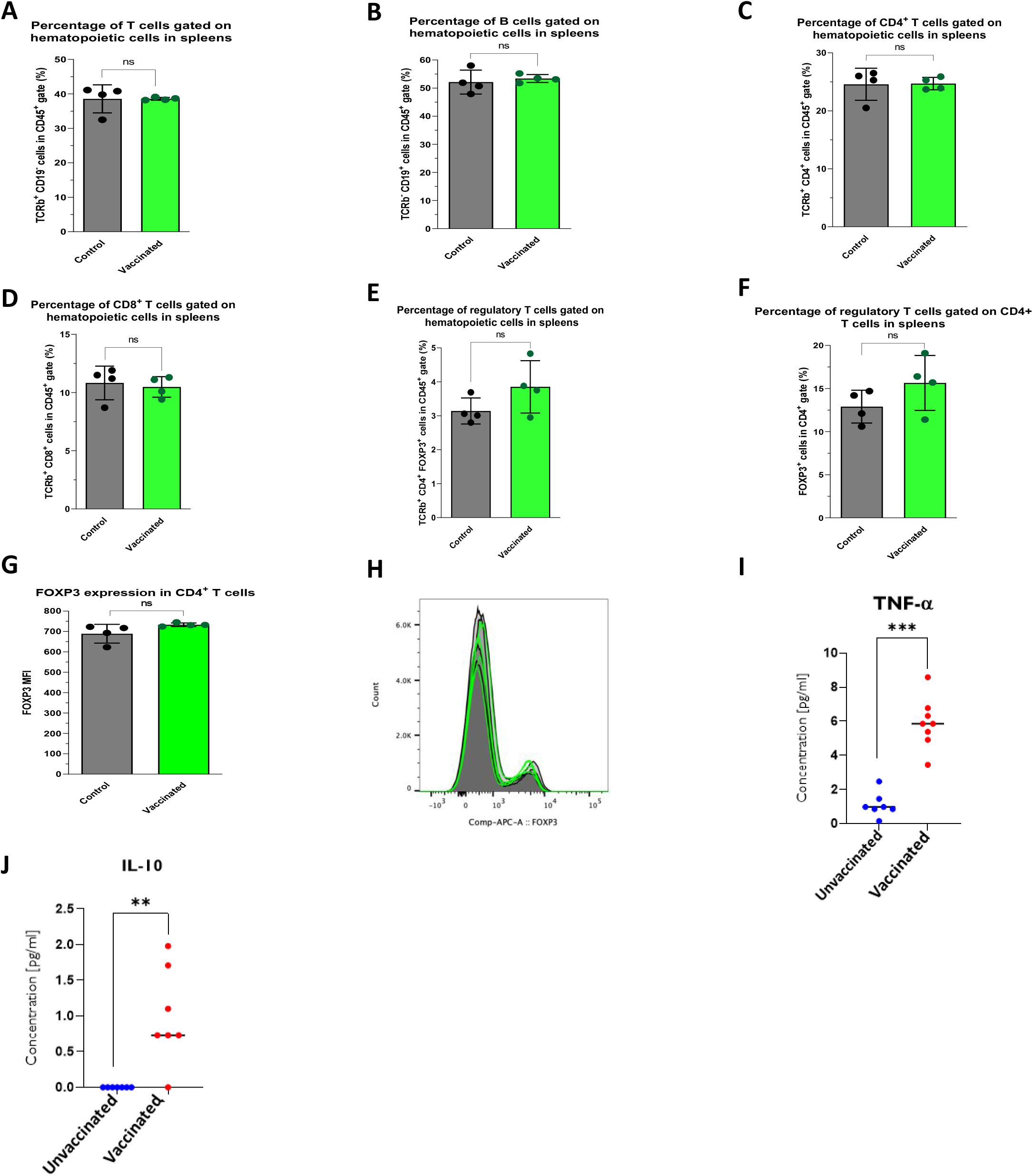
Evaluation of the cellular response after immunization by *E. coli* E11 ΔaroA Comparison of the expression of T cells (**A**), B cells (**B**), CD4+ T cells (**C**), CD8+ T cells (**D**) and regulatory T cells (**E**) in the haematopoietic population of splenocytes in the group of vaccinated and non vaccinated mice. No significant differences were observed for these different cell populations between the two groups (P>0.05). Comparison of regulatory T cell expression in the DC4+ T cell population (**F**) and FOXP3 expression in the DC4+ T cell population (**G**). A trend of increased regulatory T was observed in the vaccinated group of mice but was significantly different between the two groups (P>0.05). (**F**) A representation of the fluorescence for FOXP3 labelling. The green lines correspond to samples from treated mice, the grey areas correspond to samples from untreated mice. An almost perfect superposition of all samples can be observed. The concentration of TNF (**I**) and IL-10 (**J**) in the sera of vaccinated mice seven days after the third dose was significantly increased, compared to those of unvaccinated mice (p<0.05). IL-2, IL4, IFN gamma and IL-17A were absent in the sera of the vaccinated and unvaccinated groups. Asterisks indicate significant differences (Student’s t test: **P value < 0.01 and ***P value < 0.001) and ns non-significant

### Evaluation of the protection of vaccinated female mice

#### In vitro protection

To evaluate the *in vitro* protection mediated by the polyclonal antibodies generated by the vaccination, bactericidal assays against three strains of *E. coli K1* (E11, S88 and RS218) were performed. We detected strong bactericidal activity in serum of mice immunized up to 1/100 dilution in the presence of 3–4-week-old rabbit baby supplement against all *E. coli* k1 tested strains. (**Figure 4A**). To confirm the specificity of the bactericidal activity, another pathogenic Gram-negative bacteria, *Pseudomonas aeruginosa* PA14, previously used in killing experiments^31^ was tested. No bactericidal activity was found against *P. aeruginosa* (**Figure 4B**).

**Figure 4:**
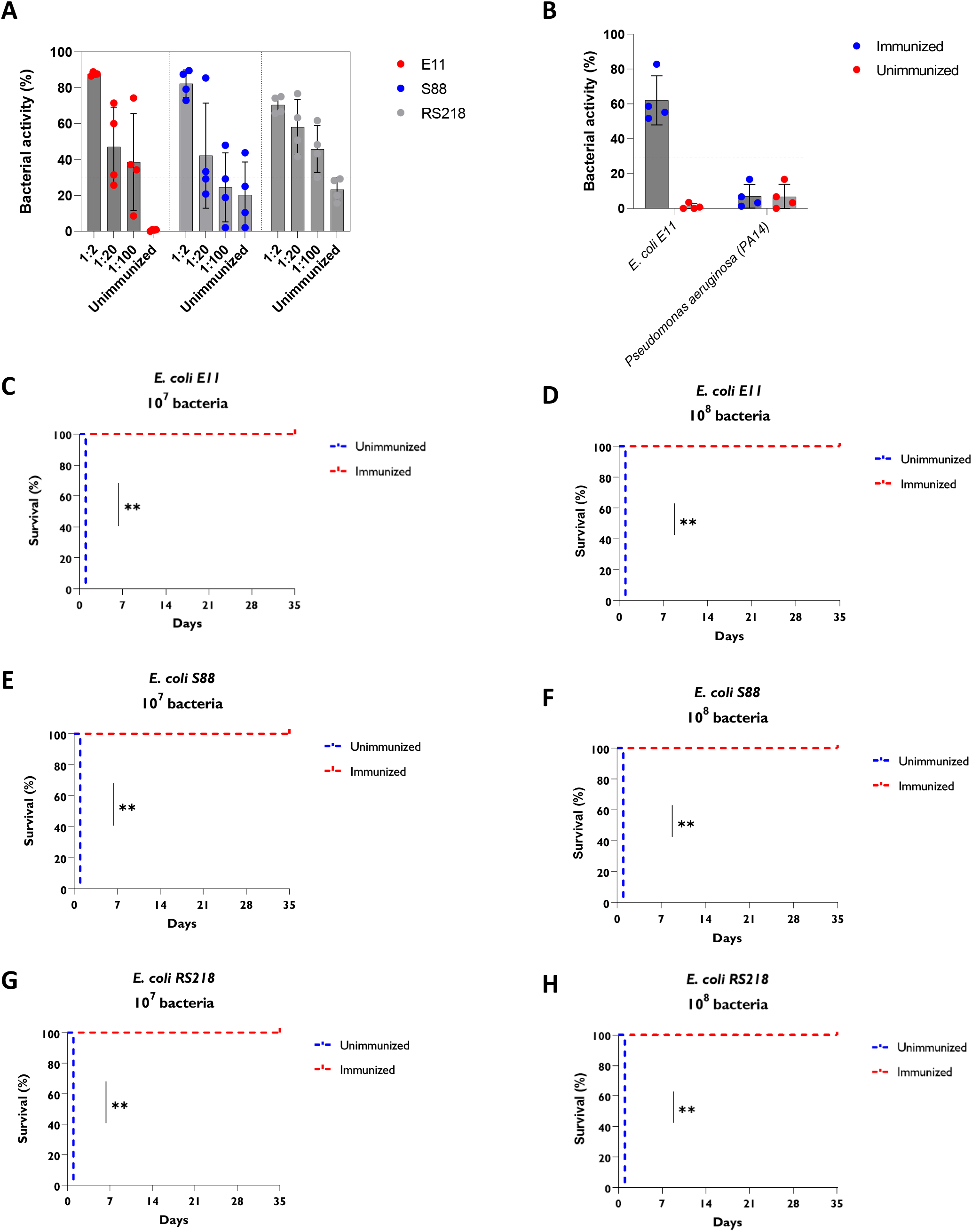
Evaluation of maternal protection by *E. coli* E11 ΔaroA vaccine **(A)** Percentage of bacterial activity obtained with different dilutions (1/2, 1/20 and 1/100) of sera from immunized and inactivated mice in the presence of control sera inactivated on the different E. coli strains E11, S88 and RS218 and using 3-4 week old rabbit baby complement. (**B**) Percentage of activity obtained with inactivated sera from mice immunized on *Pseudomonas aeruginosa* PA14 and using 3-4 week old rabbit baby complement Bactericidal activity is considered potentially significant in vivo if in vitro killing is greater than 30%.On day 7 post-vaccination, groups of vaccinated female Balb/C mice (n=4 per group) were challenged with lethal doses of 107 (**C**) and 108 (**D**) of *E. coli* E11 WT, 107 (**E**) 108 (E) of *E. coli* S88, and 107 (**F**) 108 (**G**) of *E. coli* RS218. Asterisks indicate significant differences (log-rank test (Mantel-cross): **P value < 0.01).

#### In vivo protection

To evaluate the protection conferred *in vivo* by our vaccine candidate, adult female mice were challenged with two lethal doses of 10^7^ and 10^8^ of three *E. coli* K1 (E11, S88 and RS218). Vaccinated mice showed significantly (p<0,01) better survival than that of the non-immune group, which showed a 100% mortality rate within 24 hours. (**Figure 4C, D, E, F, G and H**). No abnormalities were observed 35 days after protection.

Collectively, these results indicate that vaccination with *E. coli* E11 Δ*aro*A generated a strong humoral response driven by polyclonal antibodies and that these antibodies can mediate the in vitro killing of *E. coli* K1 and protect in vivo vaccinated animal against severe infection caused by *E. coli* K1.

### Evaluation of maternal antibodies in mouse neonates

To assess the vertical transfer of maternal antibodies to mouse pups, female mice were immunized with three doses of our vaccine and then mated with male mice of the same Balb/C genetic background (**Figure 5A**). Seven days *post-partum*, the pups were sacrificed, blood was collected and antibody titers specific to *E. coli* K1 quantified. We found antibodies able to bind to several strains of *E. coli* K1 in the sera of neonates born from immunized mothers but not in the control group (**Figure 5B**). The maternal antibodies found in neonates were confirmed to be polyclonal by Western Bots (**Figure 5C**). They were absent in the sera of neonates born from non-immunized mothers (**Figure 5D**).

**Figure 5:**
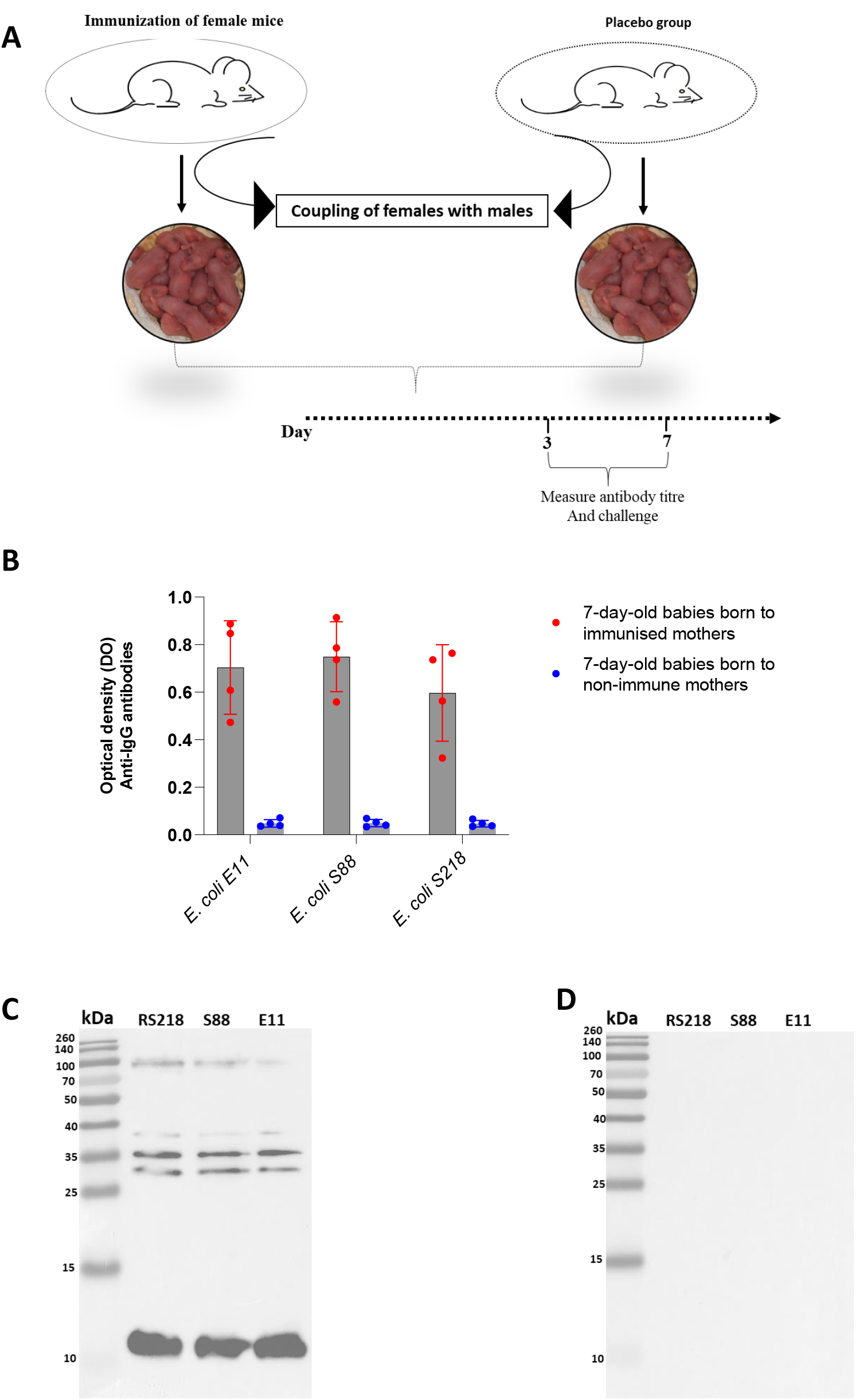
Evaluation of the presence of maternal specific antibody in mouse pups (**A**) 7 days after the last vaccination dose, vaccinated and unvaccinated (placebo) Balb/C female mice were mated with Balb/C males, after gestation and delivery the baby mice groups of 3 and 7 day old mice from the immunised and non-immunised dams were sacrificed for sera were collected for evaluation of maternal antibody titers specific to *E. coli* E11, S88 and RS218 (**B**) by whole cell ELISA and for the detection of polyclonal antibodies by western blot (**C**) and the presence of the control group (**D**) (serums of the baby mice from non-immunised mothers).

All these results confirm the vertical transfer of maternal antibodies to the neonates.

### The effect of maternal vaccination on the protection of the neonates

The protection of pups born from vaccinated mother was evaluated for early-set and late-set neonatal infections.

#### Early-set infection

Neonates of 3 days of age were challenged with 10^4^ and 10^5^ of *E. coli* K1 S88 (**Figure 6A and 6B**) as well as 10^5^ of *E. coli* RS218 (**Figure 6C**) and 10^5^ of *E. coli* E11(**Figure 6D**). A very strong protection (p=0,0082) was observed with 100% mortality rate observed in pups born from unvaccinated mother and 100% survival for neonates born from vaccinated mothers.

**Figure 6:**
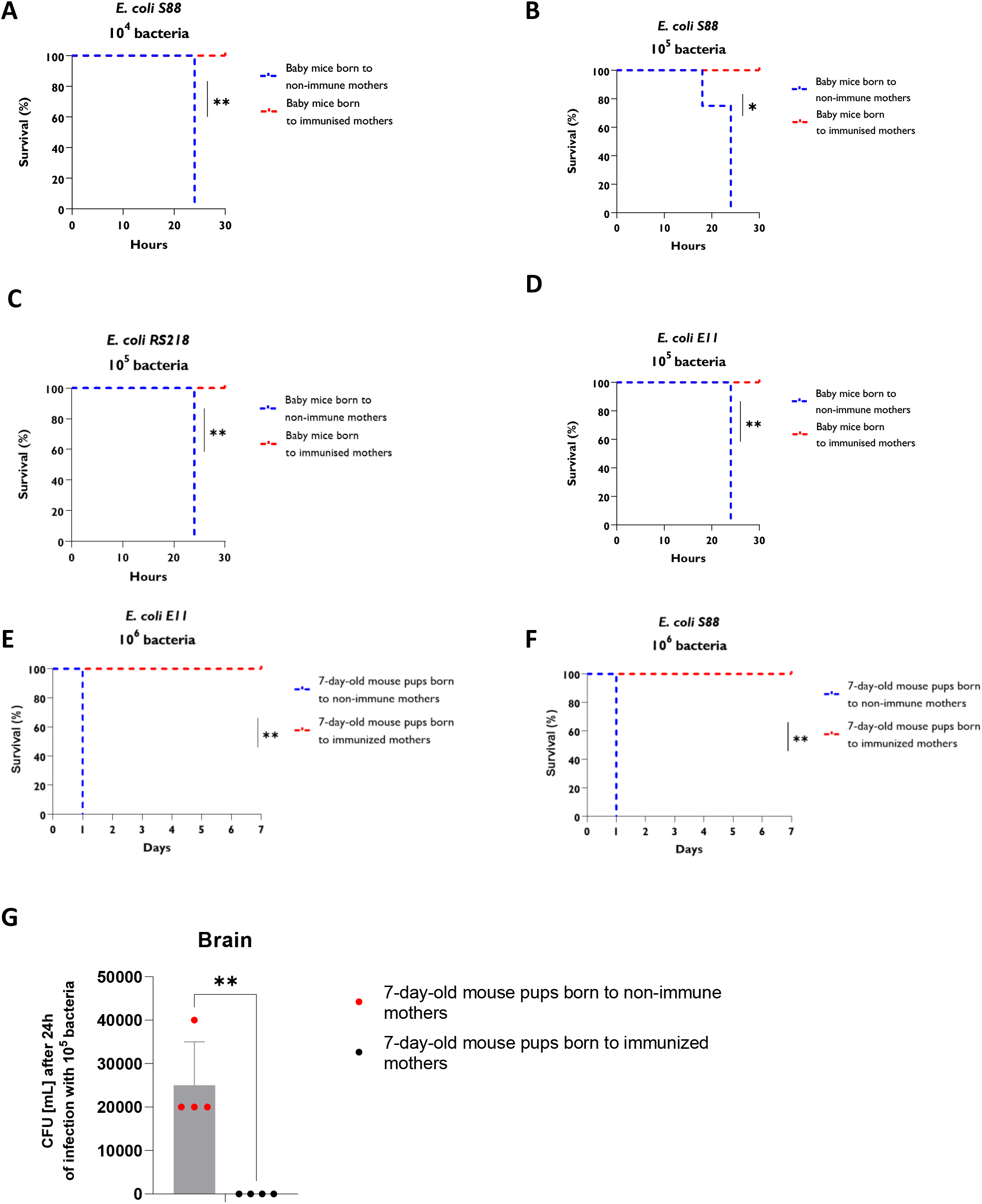
Maternal immunization with *E. coli* K1 ΔaroA against neonatal meningitis Three days after delivery, groups of four born vaccinated and unvaccinated female mice were injected intraperitoneally with CFU’s of 104 (A) and 105 (B) of *E. coli* S88 ; 105 of RS218 (C) and 105 of E11 WT (D). Seven days after delivery, groups of 4 babies mice born to vaccinated and unvaccinated females were also given CFU’s of 106 *E. coli* E11 WT (E) and 106 E. coli S88 (F). Protection was assessed for 7 days. (G) CFU count of the brain 24 hours after infection with 106 CFU of *E. coli* E11 WT bacteria in three-day-old mouse pups. Asterisks indicate significant differences (log-rank test (Mantel-cross) : **P-value =0.008 and *P-value =0.01).

#### Late-set infection

A 100% protection was also observed in 7-day-old pups born to immunized mothers after challenge with 10^6^ of *E. coli* k1 E11 and S88 strains (**Figure 6E et 6F**).

While we previously reported a high level of bacteria in the brains of the baby mice after IP infection by *E. coli* K1 E11^23^, no bacteria was found in the brain of the 7 days old pups born from immunized mothers (**Figure 6G**)

Collectively, these results indicate that maternal antibodies transferred to the newborn can effectively protect against severe infection caused by different *E. coli* K1 strains against both early and late-onset infections.

## Discussion

Antibiotic-resistant *E. coli* strains have been found in premature infants and in women who gave birth prematurely^32,33^. The efficacy of advanced antibiotic therapy for *E. coli* K1 meningitis is limited by the emergence of antibiotic resistance by some *E. coli* K1 strains^16,34,35^. Even when caused by strains susceptible to antibiotics, there is a high rate of mortality and sequalae^36^ after treatment of *E. coli* K1 meningitis. In face of these ongoing issues, vaccines could be a rapid and effective alternative solution. The efficacy of a vaccine is driven by its safety and its immunogenicity. It is known that attenuated vaccines induce rapid, long-lasting and effective immunity^17,37–39^. While not recommended during pregnancy, they can be considered in women with a pregnancy project, before conception, to generate an immune response that could be later transferred to the neonates. The results of our study indicate that the *aor*A-deleted *E. coli* K1 E11 strain is a good candidate for an attenuated vaccine with high immunogenicity and very good preclincal in vitro and in vivo protection. We selected Δ*aro E. coli* K1 to develop our attenuated vaccine candidate because (i) *aro*A was found to be a virulence factor for *E. coli* K1 in our previous work and (ii) because this approach was successful with other pathogens^25,26^.

The specific immunity of newborns to pathogens relies mainly on antibodies of maternal origin^40^. Maternal antibodies are most needed for offspring protection in contrast to maternal T cells due to tissue antigen (HLA in particular) differences between mother and fetus^41^. In this study, we show that vaccination with the Δ*aro E. coli* K1 induces a very strong humoral immunity with very high bactericidal activity. Our data also show that the antibodies induced by immunization with this vaccine candidate were sufficient for 100% protection of female mice against various lethal doses of the *E. coli* K1 strains. Studies have shown that maternal antibodies are transferred to their offspring via the placenta and breast milk^42^ and that these maternal antibodies alone are required for protection against perinatal infections^16^.

In this study, we showed the presence of maternal polyclonal antibodies specific to *E. coli K*1 in pups born to immunized mothers. This antibody response was specific to *E. coli* K1. The female mice were vaccinated prior to mating, demonstrating the long-lasting efficacy of the immunity protection induced by the *aorA* mutant *E. coli* K1 vaccine candidate.

In addition, our data also indicate effective protection of pups born from vaccinated mothers. This protection effective against both early-set^43^ and late-set^44,45^ infections, against different virulent *E. coli* strains causing meningitis and at different lethal doses. Furthermore, no bacteria were detected in the brains of puppies born from vaccinated mothers. No detectable T cell response was found in vaccinated mothers, but this had impact on the neonates protection.

Maternal antibodies can access fetal neural tissue through the developing blood-brain and blood-nervous barrier and prevent infection by vertically transmitted pathogens^32^. One of the pathogenic powers of *E. coli* K1 is their ability to cross the blood-meningeal barrier and to multiply in cerebrospinal fluid^46^. Therefore, the presence of maternal antibodies in the sensitive tissues of newborns could confer a very strong protection.

## Conclusion

In this study, we present an attenuated *E. coli* K1 vaccine candidate by deletion of the *aroA* gene that induces strong humoral immunity capable of successfully killing *E. coli* K1 *in vitro* and successfully protecting adult mice *in vivo*. We also showed the presence of maternally derived polyclonal antibodies in pups born to mothers immunized with our vaccine candidate and full protection of these pups against different strains of *E. coli* K1 *in vivo*. As there is a very high risk of a second preterm birth in women with a first preterm birth and an increased risk of neonatal *E. coli* K1 meningitis in future newborns born to these women, our study provides a strong pre-clinical proof-of-concept that an attenuated vaccine might be good approach to be further developed for this targeted population, specially when planning a future pregnancy.

## Methods

### Ethical Statement

The mouse studies were conducted in accordance with the recommendations of the French University and Research Institution Standards for the Use of Laboratory Animals. The protocols were validated by the ethical committee for animal use (Folder number: 2020011519022360; approval number A75-14-08). Intraperitoneal injections were performed under Ketamine and zylazine induced anesthesia. Every effort was made to minimize animal suffering.

### Bacteria and cells

The different *E. coli K1* strains were provided by Professor Stéphane Bonacorci, head of the French national *Escherichia coli K1* centre (**Table 2**). The *Hela* cell line was provided by HeLa CRM-CCL-2™ and maintained at 37°C with 5% CO2 in Dulbecco’s high glucose modified Eagle’s medium (DMEM, Invitrogen, Carlsbad, CA) with 10% fetal bovine serum (FBS, Dominique Dutscher, France).

### Construction of the aroA mutant

The *aroA* gene is present in *Escherichia coli* strains *E. coli* E11 and consists of 1284 nucleotides. The methodology we used to deletion the *aroA* gene in *Escherichia coli* was described by Datsenko in the 2000s^28^. This technique use recombination between homologous sequences in the vicinity of the gene to be deleted and a previously generated DNA cassette. Recombination is facilitated by the Reda/Redb proteins of the pRed plasmid. Primers containing homology sequences (20 nucleotides) with the kanamycin resistance cassette (1737 bp) contained in the genomic DNA of *E. coli* E11 *∆csga* that we wish to amplify were used. These primers also contain sequences homologous adjacent (50 nucleotides) to the *aroA* gene of *E. coli* E11(**Table 1)**. PCR was performed to generate a DNA product containing the kanamycin resistance cassette measuring 1.7kb which was subsequently column purified. The pRed plasmid was transferred into our *Esherichia coli* strains by electroporation technique. After electroporation, the strain was grown overnight on solid medium (LB) containing 100 µg/ml ticarcillin in order to select the transformed bacteria containing the pRed plasmid. The resulting colonies were then re-isolated onto LB agar plates containing 100 µg/ml ticarcillin and tested by PCR with primers homologous to the ampicillin resistance gene contained in pRed. The proteins required for homologous recombination were induced by culturing Escherichia coli plus pREd with L-arabinose. Then, the kanamycin resistance cassette containing the homologous sequences adjacent to the *aroA* gene was transferred into *Escherichia coli* plus pRed by electroporation (the pRed plasmid is lost during this step by culture on liquid medium at 37°C). At the end of this step, the bacteria are grown overnight on LB medium containing 20, 50 and 100 µg/ml kanamycin in order to select mutants that have inserted the kanamycin resistance cassette and deleted their *aroA* gene by homologous recombination.

### Bacterial growth curve

Wild type and *∆aroA Escherichia coli* strains were grown for 48h at 37°C on LB medium with or without the addition of 40µg/ml tryptophan, 40µg/ml tyrosine, 40µg/ml phenylalanine. During this period, bacterial growth was measured by Optical Density. Subsequently, wild type and *∆aroA Escherichia coli* E11 strains will be studied using the same protocol.

### Determination of virulence of the wild-type and ΔaroA mutant strains

To assess virulence, six-week-old female mice were used to determine the lethal dose of the *E. coli E11* wild-type and ΔaroA mutant strains. For each strain, 60 mice were divided into 5 groups (n = 4 mice each). The bacterial strains were then grown to an OD600 value of 0.6-0.8, washed twice with sterile PBS, and resuspended in NaCl. Mice in each group were intraperitoneally injected with 10^9^, 10^8^, 10^7^, 10^6^ and 10^5,^ CFU of bacteria diluted in 200 µl of NaCl, respectively. Mouse mortality was observed for one week.

### Bacterial adhesion assays on epithelial cells and macrophages

The two strains of bacteria *E. coli E11* (WT and *ΔaroA*) were cultured in LB at 37°C overnight. The next day, a preculture was started with an OD=0.05 and allowed to grow until an exponential culture OD600 of between 0.6-0.7 was reached. Then, the strains were pelleted by centrifugation at 5000×g for 10 minutes at 4°C and the resulting supernatant was discarded and the pellet was washed twice with PBS and resuspended in Eagle’s modified medium. Dulbecco (DMEM) without antibiotics. *Hela* epithelial cells and Row 264.7 macrophages were grown in 24-well plates at 37°C in a 5% CO2 humidified atmosphere for infection assays. The cells were washed and incubated with the bacteria containing in the medium with an infection triplicate. The infected cells were placed at 37°C in a humidified atmosphere at 5% CO2. For the adhesion plate, after one hour of infection the supernatants were harvested and stored at -20°C. After being washed five times with PBS to eliminate non-adherent bacteria, the adherent cells of the plates intended for adhesion were lysed with 500 μL of Triton X-100 at 0.2% for 10 min on ice. The cell suspension was then serially diluted 10-fold with PBS and plated on LB agar plates to determine adherence frequency. In order to quantify the cytokines, a plate of each cell after 1 h of adhesion and washing was then recultured with medium containing 100 μg/mL of gentamicin and incubated at 37°C for 3 h and the cell supernatants were were collected and stored at -20°C for cytokine assays.

### Antibody response from Balb/C femelle mice

To assess the titer of *E. coli*-specific antibodies, six-week-old female mice were dosed three times with the Δ*aroA* mutant strain of *E. coli E11* in the presence of a control group that received placebo (NaCl). Blood samples were taken from the retroorbital sinus seven days after each dose and up to 28 days after the last dose. Antibody titres to the different strains of E. coli K1 and non-K1 causing neonatal meningitis were assessed by a whole cell ELISA. Whole cell ELISAs were performed by standard methods^47^. To assess heat-stable antigens, bacteria were inactivated in a water bath at 70°C for 60 min in PBS, then centrifuged for 10 min at 20,000 × g in a microcentrifuge. The pellet was resuspended in 0.01 M sodium phosphate buffer (pH=7.0) and 100 µl of this suspension at OD=0.1/mL was used to coat microtitre plates for ELISA. The anti-mouse anti-sp (Rabbit anti-mouse IgG H&L (ab6709) coupled to HRP diluted 1:10000 was used. He histogram of antibody titres was plotted using GraphPad Prism 9 software (La Jolla, CA).

### Western blot essay

Bacterial cells from the three *E. coli K1* strains (E11, S88 and RS218) were collected by centrifugation of a 1 mL bacterial culture grown at 37 °C in OD600 of 1.00. The bacterial cells were then suspended in sample lysis buffer RIPA. Lysed samples were centrifuged and 10 μL of the supernatant was electrophoresed in a 4-15% SDS-Polyacrylamide gel (SDS-PAGE, BioRad). Proteins from gels not stained with Coomassie Blue were transferred using the transfer system to nitrocellulose membranes, which were probed using the BioRad Western Transfer System according to the manufacturer’s instructions. Mouse sera were pooled according to immunised and non-immunised group and a rabbit anti-mouse HRP peroxidase antibody (diluted 1:5000 in sterile water) was used for detection. The chemiluminescent substrate (ECL Rev-elBlot) was used for development.

### Spleen cell phenotype analysis

After 7 days immunisation, spleens from BALB/c mice were harvested for lymphocyte analysis by flow cytometry. Red blood cells were removed from splenic cell suspensions by lysis with ammonium chloride buffer. Cells were then incubated with Live/Dead Aqua kit (Invitrogen L34957) in RT PBS according to the manufacturers recommendations. Then cells were surface stained with anti-CD45 PerCP-Cy5.5 (BioLegend 103132), anti-CD19 APCeF780 (eBiosci-ence 47-0193-80), anti-TCRb AF700 (BioLegend 109224), anti-CD8 BV605 (BD Horizon 563152), and anti-CD4 PB (100534) in cold PBS containing 0.5% BSA. Cells were then fixed and permeabilized with True-Nuclear™ Transcription Factor Buffer Set (BioLegend 424401) following the manufacturers instructions and were then stained with anti-FOXP3 APC (Invi-trogen 17-5773-82) diluted in permeabilization buffer. Cells were then washed and resuspended in cold PBS containing 2% FCS. 10^5 stained cells were analyzed on a Fortessa flow cytometer (BD Biosciences, San Jose, CA). Doublets were excluded from the analysis by using appropri-ate FSC/SSC gates. Live cells were gated as Live/dead negative cells. Then each immune sub-population was characterized as follows: B cells as CD45^+ TCRb^-CD19^+, T cells as CD45^+ CD19^-TCRb^+, CD4 T cells as CD45^+ CD19^-TCRb^+ CD8^-CD4^+, CD8 T cells as CD45^+ CD19^-TCRb^+ CD4^-CD8^+, regulatory T cells as CD45^+ CD19^-TCRb^+ CD8^-CD4^+ FOXP3^+. Data were analyzed with Flowjo software, results are ex-pressed as the percentage of the selected population out of all hematopoietic cells.

### Quantification of cytokines

The production of human cytokines IL-10 or IFN-gamma, TNF-alpha, IL-4, IL-6 and TGF-beta1 was quantified in serum from immunized and non-immunized mice using the CBA (cy-tometric beads array -BD™) method, according to the manufacturer’s protocols. CBA param-eters were acquired by the BD Accuri cytometer and analysed using HEC software. All assays were expressed as pg/mL of each cytokine, determined by a standard curve of cytokine concen-trations.

### Bactericidal activity assay (BSA)

Serum bactericidal assays (BSA) against the three majority strains of *E. coli K1* (E11, S88, RS218) following the standard BSA protocol^48,49^. The different strains tested were grown at 37 °C in exponential phase from an overnight culture in LB medium for two hours and washed twice in PBS. 10 μl of washed bacteria were resuspended in PBS, without additional Mg2+ or Ca2+, was added to 10 μl of serum, 10 μl of three-to four-week-old rabbit baby complement, and 10 μl of PSS in a 96-well plate to obtain a final concentration of 10^3^ CFU/mL in serum, and incubated at 37°C. Viable amounts of the different *E. coli K1* strains were determined for the bacterial inoculum after 60 minutes of exposure to serum by serial dilution in PBS, followed by overnight growth at 37°C on LB agar. *E. coli K1* killing was calculated by comparing the concentration of viable *E. coli K1* at the beginning of the assay with the concentration of viable bacteria at each time point. Serum bactericidal assays were performed with serum from heat-inactivated mice as a negative control. Sera that showed bactericidal activity (>30%) in two or more dilutions were considered positive.

### Evaluation of the protection of the attenuated E. coli E11 ΔaroA vaccine

To evaluate the protection of the attenuated *E. coli E11* Δ*aroA* vaccine, groups of six-week-old female Balb/C mice were immunized with doses of live *E. coli E11 ΔaroA* vaccine at a rate of one dose per week and each dose contained 10^5^ CFU of bacteria diluted in 200 µl NaCl. One week after the third dose, the mouse groups were given lethal doses (10^7^, 10^8^ CFU) of the three major strains responsible for neonatal meningitis, namely *E. coli E11, S88* and *RS218*. Protec-tion was assessed for 35 days.

### Evaluation of maternal vaccine antibody transfer and protection of infant mice born to vaccinated mothers

To assess the protection of vaccinated maternal pups, groups of six-week-old female Balb/C mice were immunized with doses of live *E. coli E11* Δ*aroA* vaccine at a rate of one dose per week and each dose contained 10^5^ CFU of bacteria diluted in 200 µl NaCl and control groups received the same volume of saline (Nacl). The different groups of female mice were matched with male mice, also six weeks old. Three and seven days after birth, 2 groups of 4 babys mice from the vaccinated and unvaccinated dams respectively were decapitated and blood was collected for quantification and detection of *E. coli K1* specific antibodies by whole cell ELISA and western blot respectively. The dams were euthanised and blood was also collected for quantification of specific antibodies. Furthermore, groups of 4 babys mice originating respectively from vaccinated and non-vaccinated mothers of 3 and 7 days of life were given different doses of the *E. Coli K1* strains E11, S88 and RS218 and the survival was evaluated.

---

**Supplementary figure 1:**
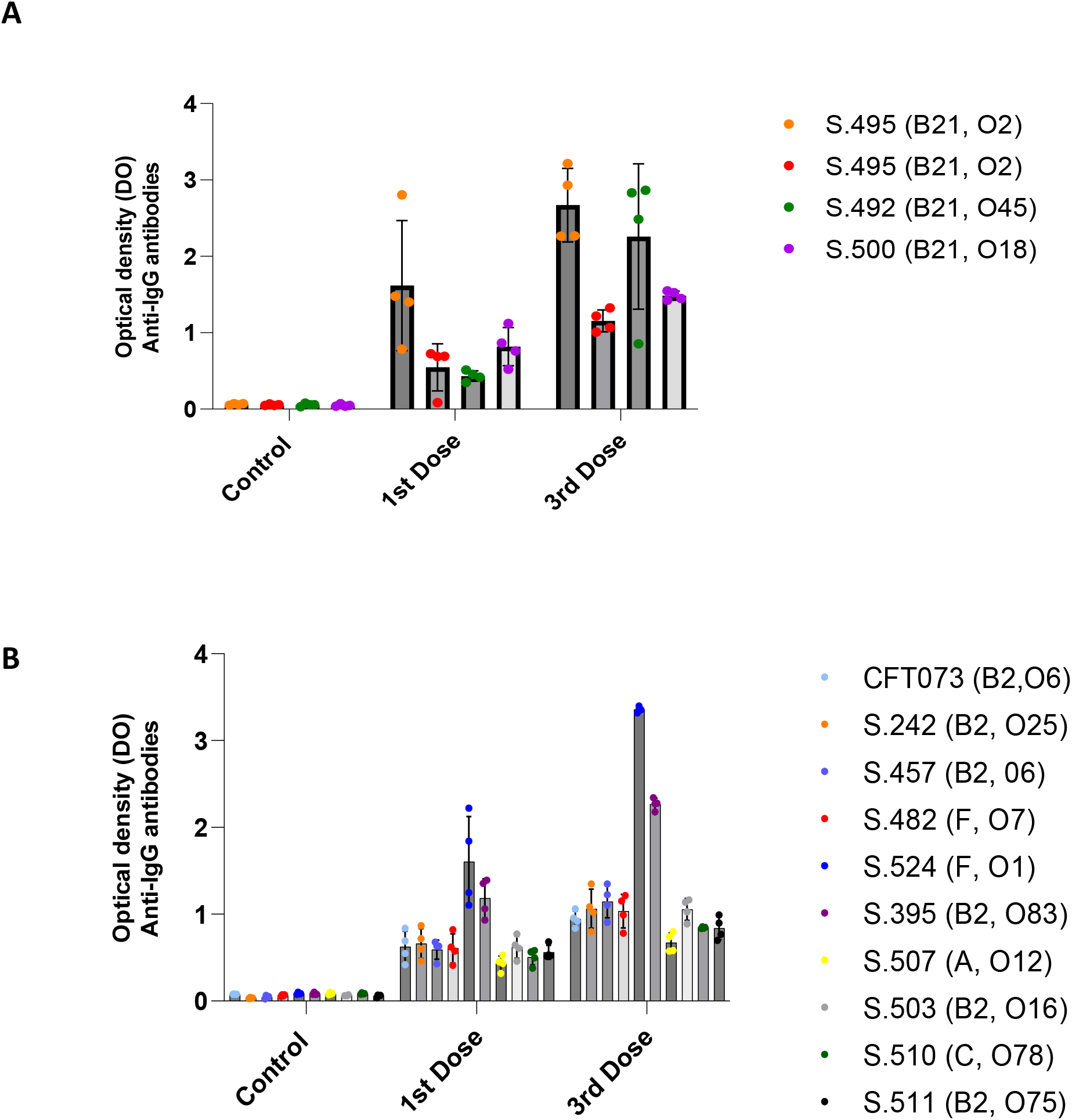
Antibody responses to other E. coli K1 and non-K1 strains responsible for neonatal meningitis in the serum of mice after immunisation with *E. coli* E11 ΔaroA (**A**) IgG response to other *E. coli K1* strains ((S.495 (B21, O2) S.492 (B21, O45) S.500 (B21, O18) S.509 (B21, O1)). (**B**) IgG response to other *E. Coli* non-K1 E. coli strains (CFT073 (B2, O6) S.242 (B2, O25) S.457 (B2, 06) S.482 (F, O7) S.524 (F, O1) S.395 (B2, O83) S.507 (A, O12) S.503 (B2, O16) S.510 (C, O78) S.511 (B2, O75)) response to neonatal meningitis IgG responses were measured using an established whole cell ELISA. Titers are presented as optical density. Sera were diluted 1/50.

## Notes

### Competing Interest Statement

The authors have declared no competing interest.

## References

1. Vogel, J. P. et al. The global epidemiology of preterm birth. Best Practice & Research Clinical Obstetrics & Gynaecology 52, 3–12 (2018).

2. Blencowe, H. et al. National, regional, and worldwide estimates of preterm birth rates in the year 2010 with time trends since 1990 for selected countries: a systematic analysis and implications. Lancet 379, 2162–2172 (2012).

3. Walani, S. R. Global burden of preterm birth. International Journal of Gynecology & Obstetrics 150, 31–33 (2020).

4. Goldenberg, R. L., Culhane, J. F., Iams, J. D. & Romero, R. Epidemiology and causes of preterm birth. Lancet 371, 75–84 (2008).

5. Romero, R. et al. The preterm parturition syndrome. BJOG 113 Suppl 3, 17–42 (2006).

6. Collins, A., Weitkamp, J.-H. & Wynn, J. L. Why are preterm newborns at increased risk of infection? Arch Dis Child Fetal Neonatal Ed 103, F391–F394 (2018).

7. Pammi, M. & Brocklehurst, P. Granulocyte transfusions for neonates with confirmed or suspected sepsis and neutropenia. Cochrane Database Syst Rev 2011, CD003956 (2011).

8. Bae, Y. M. et al. Effect of exogenous surfactant therapy on levels of pulmonary surfactant proteins A and D in preterm infants with respiratory distress syndrome. J Perinat Med 37, 561–564 (2009).

9. McGreal, E. P., Hearne, K. & Spiller, O. B. Off to a slow start: under-development of the complement system in term newborns is more substantial following premature birth. Immunobiology 217, 176–186 (2012).

10. Stoll, B. J. et al. Early-Onset Neonatal Sepsis 2015 to 2017, the Rise of Escherichia coli, and the Need for Novel Prevention Strategies. JAMA Pediatr 174, e200593 (2020).

11. Dawson, K. G., Emerson, J. C. & Burns, J. L. Fifteen years of experience with bacterial meningitis. Pediatr Infect Dis J 18, 816–822 (1999).

12. Gaschignard, J., Levy, C., Bingen, E. & Cohen, R. Épidémiologie des méningites néonatales à Escherichia coli. Archives de Pédiatrie 19, S129–S134 (2012).

13. Usui, R. et al. Vaginal lactobacilli and preterm birth. 30, 458–466 (2002).

14. Krohn, M. A., Thwin, S. S., Rabe, L. K., Brown, Z. & Hillier, S. L. Vaginal Colonization by Escherichia coli as a Risk Factor for Very Low Birth Weight Delivery and Other Perinatal Complications. The Journal of Infectious Diseases 175, 606–610 (1997).

15. Gu, H. et al. Rational Design and Evaluation of an Artificial Escherichia coli K1 Protein Vaccine Candidate Based on the Structure of OmpA. Front Cell Infect Microbiol 8, 172 (2018).

16. Zhang, J. et al. Development of a chitosan-modified PLGA nanoparticle vaccine for protection against Escherichia coli K1 caused meningitis in mice. J Nanobiotechnology 19, 69 (2021).

17. Patel, C. D. et al. Maternal immunization confers protection against neonatal herpes simplex mortality and behavioral morbidity. Sci Transl Med 11, eaau6039 (2019).

18. Shan, C. et al. Maternal vaccination and protective immunity against Zika virus vertical transmission. Nat Commun 10, 5677 (2019).

19. Locht, C. & Mielcarek, N. Live attenuated vaccines against pertussis. Expert Review of Vaccines 13, 1147–1158 (2014).

20. Gruslin, A. et al. Immunization in pregnancy. J Obstet Gynaecol Can 31, 1085–1101 (2009).

21. MPH, I. T. G., MD. Vaccines for women: Before conception, during pregnancy, and after a birth. Harvard Health https://www.health.harvard.edu/blog/vaccines-for-women-before-conception-during-pregnancy-and-after-a-birth-2020011018649 (2020).

22. Vaccines for Women Before Pregnancy, During Pregnancy and After Childbirth. During Pregnancy https://www.health.gov.il/English/Topics/Pregnancy/during/Pages/vaccine_pregnant.aspx.

23. Pons, S. et al. A high-throughput sequencing approach identifies immunotherapeutic targets for bacterial meningitis in neonates. 2022.12.22.521560 Preprint at https://doi.org/10.1101/2022.12.22.521560 (2022).

24. Xing, X.-J. et al. Functional characterization of 5-enopyruvylshikimate-3-phosphate synthase from Alkaliphilus metalliredigens in transgenic Arabidopsis. J Microbiol Biotechnol 24, 1421–1426 (2014).

25. Felgner, S. et al. aroA-Deficient Salmonella enterica Serovar Typhimurium Is More Than a Metabolically Attenuated Mutant. mBio 7, e01220–16 (2016).

26. Priebe, G. P. et al. Construction and characterization of a live, attenuated aroA deletion mutant of Pseudomonas aeruginosa as a candidate intranasal vaccine. Infect Immun 70, 1507–1517 (2002).

27. Galal, H. M., Abdrabou, M. I., Faraag, A. H. I., Mah, C. K. & Tawfek, A. M. Evaluation of commercially available aroA delated gene E. coli O78 vaccine in commercial broiler chickens under Middle East simulating field conditions. Sci Rep 11, 1938 (2021).

28. Datsenko, K. A. & Wanner, B. L. One-step inactivation of chromosomal genes in Escherichia coli K-12 using PCR products. Proceedings of the National Academy of Sciences 97, 6640–6645 (2000).

29. Lemaître, C., Bidet, P., Bingen, E. & Bonacorsi, S. Transcriptional analysis of the Escherichia coli ColV-Ia plasmid pS88 during growth in human serum and urine. BMC Microbiol 12, 115 (2012).

30. Wijetunge, D. S. S. et al. Complete nucleotide sequence of pRS218, a large virulence plasmid, that augments pathogenic potential of meningitis-associated Escherichia coli strain RS218. BMC Microbiol 14, 203 (2014).

31. Skurnik, D. et al. A Comprehensive Analysis of In Vitro and In Vivo Genetic Fitness of Pseudomonas aeruginosa Using High-Throughput Sequencing of Transposon Libraries. PLOS Pathogens 9, e1003582 (2013).

32. Zhu, M. et al. Multi-Drug Resistant Escherichia coli Causing Early-Onset Neonatal Sepsis – a Single Center Experience from China. Infect Drug Resist 12, 3695–3702 (2019).

33. Li, Y. Y., Kong, C. W. & To, W. W. K. Pathogens in preterm prelabour rupture of membranes and erythromycin for antibiotic prophylaxis: a retrospective analysis. Hong Kong Med J 25, 287–294 (2019).

34. Yamamoto, T. et al. Electron microscopic structures, serum resistance, and plasmid restructuring of New Delhi metallo-β-lactamase-1 (NDM-1)-producing ST42 Klebsiella pneumoniae emerging in Japan. J Infect Chemother 19, 118–127 (2013).

35. Blanco, J. et al. National survey of Escherichia coli causing extraintestinal infections reveals the spread of drug-resistant clonal groups O25b:H4-B2-ST131, O15:H1-D-ST393 and CGA-D-ST69 with high virulence gene content in Spain. (2011) doi:10.1093/jac/dkr235.

36. Liu, Y. et al. Escherichia coli Causing Neonatal Meningitis During 2001–2020: A Study in Eastern China. Int J Gen Med 14, 3007–3016 (2021).

37. Shan, C. et al. Maternal vaccination and protective immunity against Zika virus vertical transmission. Nat Commun 10, 5677 (2019).

38. Shan, C. et al. A single-dose live-attenuated vaccine prevents Zika virus pregnancy transmission and testis damage. Nat Commun 8, 676 (2017).

39. Griffin, D. E. Measles Vaccine. Viral Immunol 31, 86–95 (2018).

40. Fouda, G. G., Martinez, D. R., Swamy, G. K. & Permar, S. R. The Impact of IgG transplacental transfer on early life immunity. Immunohorizons 2, 14–25 (2018).

41. Zinkernagel, R. M. Maternal Antibodies, Childhood Infections, and Autoimmune Diseases. New England Journal of Medicine 345, 1331–1335 (2001).

42. Hurley, W. L. & Theil, P. K. Perspectives on immunoglobulins in colostrum and milk. Nutrients 3, 442–474 (2011).

43. Mendoza-Palomar, N. et al. Escherichia coli early-onset sepsis: trends over two decades. Eur J Pediatr 176, 1227–1234 (2017).

44. Xu, M. et al. Etiology and Clinical Features of Full-Term Neonatal Bacterial Meningitis: A Multicenter Retrospective Cohort Study. Frontiers in Pediatrics 7, (2019).

45. Basmaci, R. et al. Escherichia Coli Meningitis Features in 325 Children From 2001 to 2013 in France. Clinical Infectious Diseases 61, 779–786 (2015).

46. Xie, Y., Kim, K. J. & Kim, K. S. Current concepts on Escherichia coli K1 translocation of the blood-brain barrier. FEMS Immunol Med Microbiol 42, 271–279 (2004).

47. Fleiszig, S. Lipopolysaccharide outer core is a ligand for corneal cell binding and ingestion of Pseudomonas aeruginosa. Investigative ophthalmology & visual science (1996).

48. Lu, X. et al. A Poly-N-Acetylglucosamine−Shiga Toxin Broad-Spectrum Conjugate Vaccine for Shiga Toxin-Producing Escherichia coli. mBio 5, e00974–14 (2014).

49. Granoff, D. M. Relative importance of complement-mediated bactericidal and opsonic activity for protection against meningococcal disease. Vaccine 27, B117 (2009).

